# *In vitro* reconstitution of SARS CoV-2 Nsp1-induced mRNA cleavage reveals the key roles of the N-terminal domain of Nsp1 and the RRM domain of eIF3g

**DOI:** 10.1101/2023.05.25.542379

**Authors:** Irina S. Abaeva, Yani Arhab, Anna Miścicka, Christopher U. T. Hellen, Tatyana V. Pestova

## Abstract

SARS CoV-2 nonstructural protein 1 (Nsp1) is the major pathogenesis factor that inhibits host translation using a dual strategy of impairing initiation and inducing endonucleolytic cleavage of cellular mRNAs. To investigate the mechanism of cleavage, we reconstituted it *in vitro* on β-globin, EMCV IRES and CrPV IRES mRNAs that use unrelated initiation mechanisms. In all instances, cleavage required Nsp1 and only canonical translational components (40S subunits and initiation factors), arguing against involvement of a putative cellular RNA endonuclease. Requirements for initiation factors differed for these mRNAs, reflecting their requirements for ribosomal attachment. Cleavage of CrPV IRES mRNA was supported by a minimal set of components consisting of 40S subunits and eIF3g’s RRM domain. The cleavage site was located in the coding region 18 nucleotides downstream from the mRNA entrance indicating that cleavage occurs on the solvent side of the 40S subunit. Mutational analysis identified a positively charged surface on Nsp1’s N-terminal domain (NTD) and a surface above the mRNA-binding channel on eIF3g’s RRM domain that contain residues essential for cleavage. These residues were required for cleavage on all three mRNAs, highlighting general roles of Nsp1-NTD and eIF3g’s RRM domain in cleavage *per se*, irrespective of the mode of ribosomal attachment.

## INTRODUCTION

Many viruses subvert cellular translation and mRNA surveillance/decay pathways to impair activation of innate immune pathways and to direct the translation apparatus to viral mRNAs (Abernathy and Glaunsinger, 2015; Burgess et al., 2022). Host shut-off of translation and the resulting inhibition of type 1 interferon (IFN) induction and signaling are important aspects of the pathogenesis of members of the alphacoronavirus (α-CoV) and betacoronavirus (β-CoV) genera of *Coronaviridae*, including the β-CoVs severe acute respiratory syndrome (SARS) CoV and SARS CoV-2 (Nakagawa and Makino, 2021; Minkoff and TenOever, 2023). Shut-off induced by coronaviruses is multifaceted and involves endonucleolytic cleavage leading to the degradation of cellular mRNAs, as well as inhibition of splicing, nuclear export of mRNA and cellular translation (Banerjee et al., 2020; Finkel et al., 2021; Nakagawa and Makino, 2021; Zhang et al., 2021).

Although several coronavirus proteins have been implicated in shut-off of translation Banerjee et al., 2020; Hsu et al., 2021; Thoms et al., 2020; Zaffagni et al., 2022), nonstructural protein 1 (Nsp1) is considered to be the major pathogenesis factor: it strongly dampens innate immune responses and its mutation or partial deletion impairs replication of α-CoVs and β-CoVs in cells with an intact IFN response (Kamitani et al., 2006; Wathelet et al., 2007; Züst et al., 2007; Narayanan et al., 2008; Shen et al., 2019; Fisher et al., 2022). SARS CoV and SARS CoV-2 Nsp1s inhibit translation using a dual strategy of impairing the initiation process and inducing the endonucleolytic cleavage and subsequent degradation of cellular mRNAs (Kamitani et al., 2006; Narayanan et al., 2008; Huang et al., 2011; Finkel et al., 2012; Lokugamage et al., 2012; Mendez et al., 2021).

Nsp1 is co-translationally cleaved from the N-terminus of the ORF1a polyprotein (Snijder et al., 2003). SARS CoV and SARS CoV-2 Nsp1s are 180 a.a. long and conserved (84% identity) whereas α-CoV Nsp1s are more variable. SARS CoV and SARS CoV-2 Nsp1s have a structurally conserved ∼120 a.a.-long N-terminal core that consists of an irregular seven-stranded β-barrel, a long flanking α-helix and a short 3_10_ helix (Almeida et al., 2007; Clark et al., 2021; Semper et al., 2021; Wang et al., 2023). SARS CoV-2 Nsp1 contains an additional 3_10_ helix and β-strand. The C-terminal region (a.a. 126-180) of SARS CoV-2 Nsp1 is unstructured (Wang et al., 2023), but when bound to the 40S ribosomal subunit, a.a. 154-180 form two short α-helices (Schubert et al., 2020; Thoms et al., 2020; Yuan et al., 2020). On the 40S subunit, the C-terminal α-helical mini-domain is inserted into the entrance portion of the mRNA-binding channel where it interferes with binding of mRNA (Schubert et al., 2020; Thoms et al., 2020; Yuan et al., 2020). The exact ribosomal position of the N-terminal domain of Nsp1 which is flexibly connected to the C-terminal mini-domain is obscure, even though it has recently been shown that it cross-links to the eIF3g subunit of eIF3 and some ribosomal proteins that reside at the mRNA entrance (Graziadei et al., 2022).

The ribosomal position of the C-terminal α-helical mini-domain accounts for the mechanism for inhibition of protein synthesis by SARS CoV and SARS CoV-2 Nsp1s, which consists of steric clashing of this domain with mRNA. In contrast, the mechanism by which Nsp1 induces endonucleolytic cleavage of cellular mRNAs that is followed by their Xrn1-mediated degradation (Kamitani et al., 2009; Huang et al., 2011; Gaglia et al., 2012; Mendez et al., 2021; Fisher et al., 2022) remains totally unknown. SARS CoV Nsp1 alone does not mediate cleavage of mRNAs, and this process depends on the presence of 40S subunits and is restricted to translationally active mRNAs (Kamitani et al., 2009; Gaglia et al., 2012). Cleavage sites map to the 5’UTR and proximal coding region of capped mRNAs and to the vicinity of the initiation codon at the 3’-border of poliovirus and encephalomyocarditis (EMCV) virus IRESs (Kamitani et al., 2009; Huang et al., 2011). Hepatitis C virus (HCV) and cricket paralysis virus (CrPV) IRESs did not undergo SARS CoV Nsp1-induced cleavage in cell-free extracts (Kamitani et al., 2009). The positions of cleavages and dependence of cleavage on the mechanism of translation initiation led to the suggestion that cleavage occurs at the stage of ribosomal attachment before establishment of codon-anticodon interaction (Nakagawa and Makino, 2021). Nsp1-induced cleavage required its ribosomal association, and consistently, cleavage was inhibited by K164A/H165A substitutions in SARS CoV Nsp1 that abrogate its ribosomal binding (Lokugamage et al., 2012). Importantly, R125A/K126A substitutions at the C-terminal border of the N-terminal core domain of SARS CoV Nsp1 (Lokugamage et al., 2012) and analogous substitutions in SARS CoV-2 Nsp1 (Mendez et al., 2020) abrogated Nsp1-induced mRNA cleavage but did not affect its ability to inhibit translation, separating the RNA cleavage and the translation inhibition functions of Nsp1. Nsp1’s lack of independent cleavage activity and of sequence or structural homology with ribonucleases (Almeida et al., 2007) led to the suggestion that it recruits a specific cellular endonuclease to a subset of ribosomal initiation complexes (Nakagawa and Makino, 2021; Kamitani et al., 2009).

Here, we investigated the mechanism of Nsp1-induced cleavage by reconstituting this process *in vitro* from individual purified translational components on three mRNAs (β-globin, EMCV IRES and CrPV IRES mRNAs) that use unrelated translation initiation mechanisms.

## RESULTS

### *In vitro* reconstitution reveals that canonical translation initiation components are sufficient for SARS CoV-2 Nsp1 to induce cleavage of 5’end-dependent mRNAs

To confirm the ability of recombinant SARS CoV-2 Nsp1 to inhibit translation initiation on 5’end-dependent mRNA, we assayed its influence on 48S complex formation on (CAA)_4_-MF β-globin mRNA comprising four 5’-terminal CAA repeats that allow efficient cap-independent initiation (Pestova and Kolupaeva et al., 2002) followed by the β-globin 5’UTR (Figure 1A), a short (MF) open reading frame, a stop codon and a 110nt-long 3’UTR. 48S complexes were reconstituted *in vitro* from individual purified translational components (40S subunits, initiation factors and Met-tRNA_iMet_) in the absence or presence of Nsp1. The position of assembled ribosomal complexes was determined by toe-printing, which involves extension by reverse transcriptase of a primer annealed to the ribosome-bound mRNA: cDNA synthesis is arrested by the leading edge of the 40S subunit yielding toe-prints +15–17 nt downstream from the P site codon. Strikingly, preincubation of 40S subunits with Nsp1 not only inhibited 48S complex formation, but also yielded prominent stops at discrete positions in the 5’UTR (Figure 1B, lanes 2 and 3). Persistence of these stops when reverse transcription was done after phenol extraction (Figure 1B, lanes 12 and 13) indicated that they corresponded to sites of mRNA cleavage rather than to specific binding of translational components. The most 5’-terminal cleavage occurred 11-12 nt. from the 5’-end and downstream cleavages were all separated by ∼6-8 nt. (summarized in Figure 1A). eIF2 was not required for cleavage (Figure 1B, lane 7), whereas 40S subunits, eIF3 and group 4 eIFs (eIF4A/eIF4G_736-1115_/eIF4B) were essential (Figure 1C). Omission of eIF4B moderately reduced but did not abrogate cleavage (Figure 1B, lane 5). eIF4B had a stronger effect when the concentration of eIF4G was reduced (Figure 1D, lanes 5 and 6), whereas eIF1 and eIF1A did not influence cleavage (Figure 1D, lanes 7-9). Incubation of pre-assembled 80S initiation complexes with Nsp1 did not result in cleavage (Figure 1B, lane 11) and also did not inhibit elongation (Figure 1E). The time course of Nsp1-induced cleavage revealed progressive accumulation of shorter mRNA products indicating that cleavage is sequential and starts from the 5’-terminal site (Figure 1F). Cleavage at downstream sites could potentially occur with or without prior dissociation of truncated mRNA fragments from ribosomal complexes.

**Figure 1.**
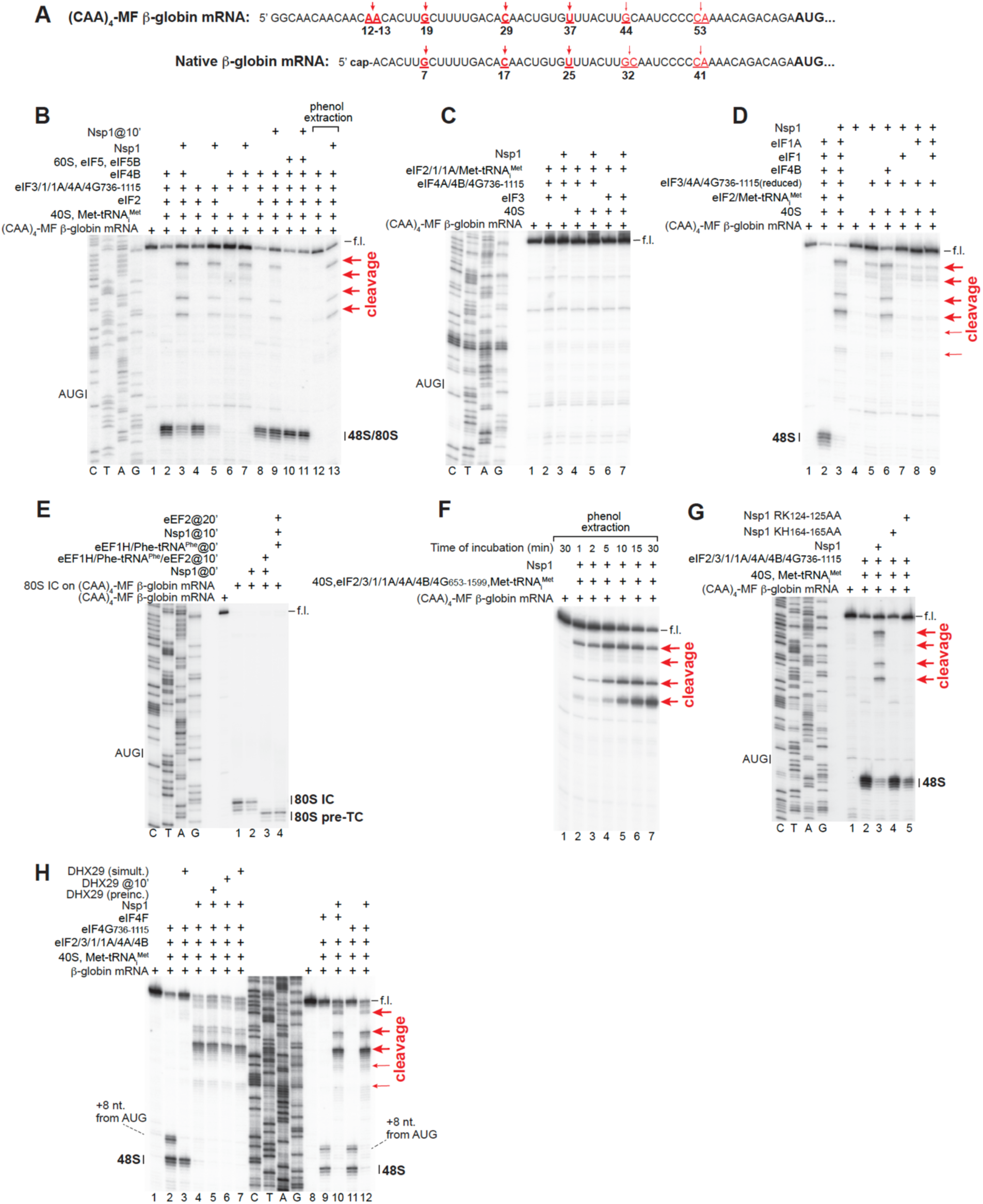
SARS CoV-2 Nsp1-induced cleavage of mRNA during 5’end-dependent initiation. (A) Schematic representation of (CAA)_4_-MF ϕ3-globin and native ϕ3-globin mRNAs, annotated to show the initiation codon (bold) and sites of Nsp1-mediated cleavage (red arrows). (B-D, F-H) Cleavage of (B-D, F, G) (CAA) _4_-MF ϕ3-globin and (H) native ϕ3-globin mRNAs depending on the presence of (B-E, G, H) *wt*, (G) RK_124-125_AA or KH_164-165_AA SARS CoV-2 Nsp1 and translational components as indicated, assayed by toe-printing, or where indicated, by primer extension by reverse transcriptase after phenol extraction of mRNA. (E) The influence of *wt* SARS CoV-2 Nsp1 on elongation by 80S initiation complexes pre-assembled on (CAA) _4_-MF ϕ3-globin mRNA, assayed by toe-printing. (B-H) Positions of the full-length (f.l.) cDNA, ribosomal complexes, and cleavage sites (red arrows) are shown on the sides of the panels. Lanes C, T, A, and G show the corresponding sequence derived using the same primer as used for primer extension.

The KH_164-165_AA Nsp1 mutant, which lacks the ability to bind to the 40S subunit (Schubert et al., 2020; Thoms et al., 2020), did not influence 48S complex formation and did not induce cleavage of (CAA)_4_-MF β-globin mRNA (Figure 1G, lane 4), whereas the RK_124-125_AA Nsp1 mutant, which inhibits translation but lacks mRNA cleavage activity *in vivo* (Mendez et al., 2020; Lapointe et al., 2021; Bujanic et al., 2022) also did not induce cleavage but retained the ability to inhibit 48S complex formation in our *in vitro* reconstituted system (Figure 1G, lane 5). Thus, the activities of Nsp1 mutants in the reconstituted system were consistent with their reported activities *in vivo*.

To determine whether the eIF4E-cap interaction influences the pattern of cleavage, we compared the effect of Nsp1 on native capped β-globin mRNA in the presence of native eIF4F or the eIF4G_736-1115_ middle domain. In the presence of eIF4F, we again observed several discrete cleavage sites with the first site slightly closer to the 5’-end than in the case of the (CAA)_4_-MF β-globin mRNA but with a similar spacing between them (Figure 1H, lane 10). The positions of cleavage sites coincided with those, starting from the second site, that were observed in the case of (CAA)_4_-MF β-globin mRNA (summarized in Figure 1A). The two most prominent cleavages (the second and third from the 5’-end) coincided with those observed in rabbit reticulocyte lysate (RRL) supplemented with SARS CoV Nsp1.^18^ Cleavage sites were identical in the presence of eIF4G_736-1115_ (Figure 1H, lanes 10 and 12). As expected, addition of DHX29 that resides at the mRNA entrance (Hashem et al., 2013a) eliminated the aberrant toe-print +8 nt. downstream from the AUG codon corresponding to incompletely closed 48S complexes (Pisareva et al., 2008) (Figure 1H, lanes 2 and 3), but did not affect cleavage irrespective of the time of DHX29 addition (Figure 1H, lanes 4-7). Thus, Nsp1-induced cleavage was not affected by the eIF4E-cap interaction or by DHX29.

Taken together, the positions of Nsp1-induced cleavage sites, the factor-dependency of cleavage and the fact that it is reduced by delayed addition of Nsp1 to pre-formed 48S complexes are consistent with the suggestion that the process occurs at the stage of ribosomal attachment before establishment of the codon-anticodon interaction (Nakagawa and Makino, 2021). However, our *in vitro* reconstitution results argue against involvement in the process of a putative, as yet-unidentified cellular RNA endonuclease.

### SARS CoV-2 Nsp1-induced cleavage of the EMCV IRES mRNA

Next, we investigated the effect of Nsp1 on the ∼450nt-long EMCV IRES (Figure 2A). Ribosomal attachment to this IRES depends on the specific interaction of its JK domain with the central eIF4A-binding domain of eIF4G (Pestova et al., 1996). The JK domain is followed by a Yn-Xm-AUG motif (Yn = pyrimidine tract; Xm = spacer) (Hellen and Wimmer, 1995), whose AUG_834_ codon is the native viral initiation codon, although low-level 48S complex formation can also occur at the nearby AUG_826_ and AUG_846_ (Figure 2B, lane 3). Again, preincubation of 40S subunits with Nsp1 resulted in the appearance of additional stops (Figure 2B, lane 4), and reverse transcription done after phenol extraction confirmed the presence of four discrete cleavage sites, ∼6-9 nt. apart and with the 5’-terminal site 2-3 nt. upstream of AUG_834_ (Figure 2B, lane 10; summarized in Figure 2A). Cleavage sites coincided with those observed in presence of SARS CoV Nsp1 in the reconstituted system (Figure 2C) and in RRL (Huang et al., 2011). Again, cleavage was 40S-dependent and was reduced by delayed addition of Nsp1 (Figure 2B, lanes 11-12).

**Figure 2.**
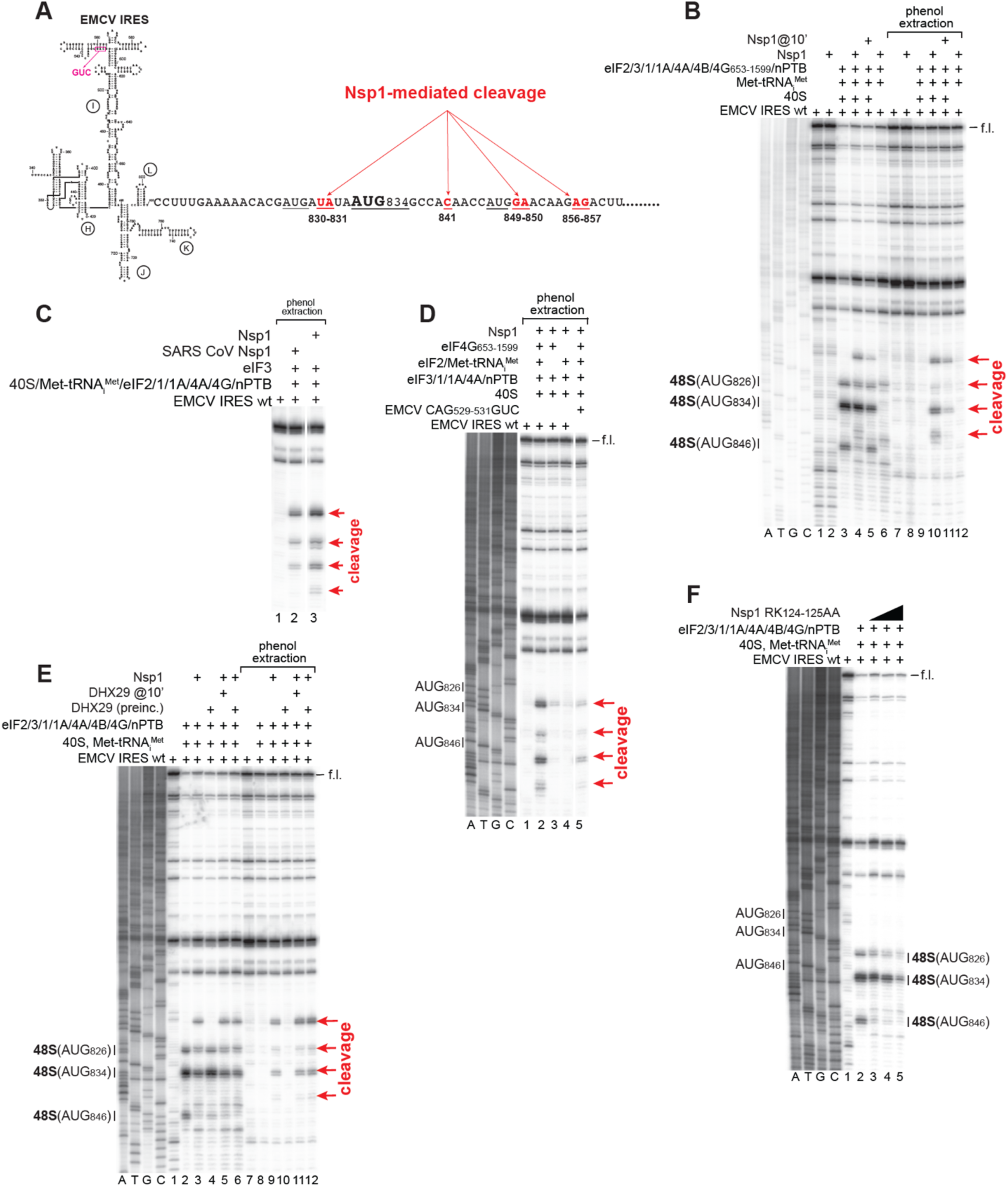
SARS CoV-2 Nsp1-induced cleavage of mRNA during EMCV IRES-mediated initiation. (A) Schematic representation of the EMCV IRES and the proximal coding region, annotated to show IRES domains H-L, the location of inactivating CAG_529-531_GUC substitutions, the initiation codon (bold) and sites of Nsp1-mediated cleavage (red arrows). (B-F) Cleavage of (B-F) *wt* and (D) CAG_529-531_GUC mutant EMCV IRES mRNA depending on the presence of (B-F) *wt* SARS CoV-2 Nsp1, (C) *wt* SARS CoV Nsp1 or (F) RK_124-125_AA SARS CoV-2 Nsp1 and translational components as indicated, assayed by toe-printing, or where indicated, by primer extension by reverse transcriptase after phenol extraction of mRNA. Positions of the full-length (f.l.) cDNA, ribosomal complexes, and cleavage sites (red arrows) are shown on the sides of the panels. Lanes C, T, A, and G show the corresponding sequence derived using the same primer as used for primer extension. Separation of lanes in panels C and D by white lines indicates that these parts were juxtaposed from the same gel.

Consistent with the essential role of eIF4G in ribosomal attachment to the EMCV IRES (Pestova et al., 1996), cleavage was abolished by omission of eIF4G (Figure 2D, lane 4). However, in contrast to 5’-end dependent mRNAs, cleavage was also strongly reduced in the absence of eIF2•GTP/Met-tRNA_iMet_ (Figure 2D, lane 3). Mutations in the IRES that influenced 48S complex formation, e.g. disruption of the secondary structure of the apex of domain I (Figure 2A), also strongly reduced cleavage (Figure 2D, lane 5). As in the case of 5’-end dependent mRNAs, cleavage was not affected by DHX29 (Figure 2E). As expected, the RK_124-125_AA Nsp1 mutant did not induce cleavage but retained the ability to inhibit 48S complex formation (Figure 2F).

In contrast to (CAA)_4_-MF β-globin mRNA, the time course of Nsp1-induced cleavage of EMCV IRES mRNA did not show accumulation of the shortest mRNA product, and the relative intensities of cleavages at the strongest first and third positions remained the same (Figure S1A). We therefore introduced various mutations downstream of the JK domain (Figure S1B) and tested their influence on Nsp1-mediated cleavage. Deletion of domain L did not influence cleavage, whereas its replacement by 4 or 8 nucleotides slightly reduced its efficiency at the last two sites (Figure S1C). Stabilization and particularly destabilization of domain L moderately reduced the efficiency of cleavage at all sites (Figure S1D, lanes 2, 4 and 6). Strikingly, introduction of three substitutions in the region surrounding the 5’-terminal site completely and specifically abolished cleavage at the first two positions (Figure S1D, lanes 2 and 8). These mutations included replacement of AUG_826_ and AUG_834_ by AUC codons. To determine whether the presence of the immediate upstream AUG codon is required for cleavages, we determined how replacing AUG_846_ by AUC and substituting the downstream AAG_853_ by AUG would influence the efficiency and the position of the two downstream cleavage sites. These changes did not influence cleavage at the two downstream sites (Figure S1E) arguing against the requirement of the immediate upstream AUG codon for Nsp1-induced cleavage. These data indicate that cleavage at the first and the second pairs of sites occur independently and suggest the possibility of some nucleotide specificity of Nsp1-induced cleavage.

In conclusion, as in the case of 5’end-dependent mRNAs, Nsp1-induced cleavage of the EMCV IRES mRNA required only canonical translational components and occurred at the stage of ribosomal attachment.

### Reconstitution of Nsp1-induced cleavage of cricket paralysis virus (CrPV) IRES mRNA reveals the minimal set of translational components that can support cleavage

Ribosomal attachment to 5’end-dependent and EMCV IRES-containing mRNAs relies on large, strongly overlapping sets of initiation factors. To investigate whether Nsp1-mediated cleavage can be achieved with a smaller set of eIFs, we took advantage of factor-independent ribosomal attachment to the CrPV IGR IRES. This IRES comprises three pseudoknots (PKI-PKIII) (Figure 3A). Stem-loops SL2.1 and SL2.3 in PKIII contain conserved apical motifs that interact with ribosomal proteins on the head of the 40S subunit (Muhs et al., 2011; Fernández et al., 2014), whereas PKI mimics the anticodon stem-loop of tRNA base-paired to a cognate codon (Costantino et al., 2008). The CrPV IRES binds directly to individual 40S subunits or to assembled 80S ribosomes (Wilson et al., 2000; Pestova et al., 2004). In the resulting complexes, the IRES is located in the intersubunit space, with PKI mimicking the tRNA/mRNA interaction in the decoding center of the ribosomal A site (Schüler et al., 2006; Fernández et al., 2014).

**Figure 3.**
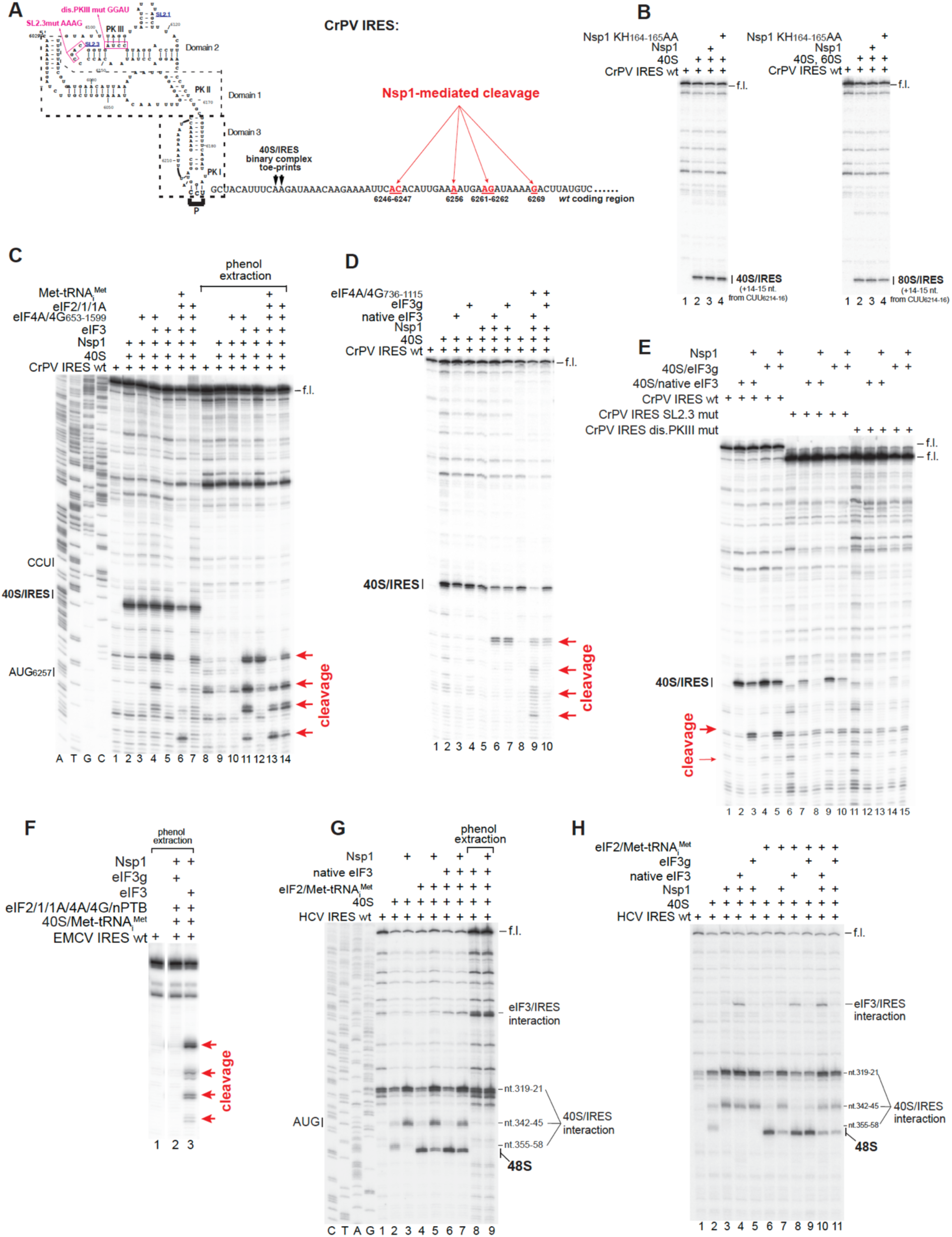
SARS CoV-2 Nsp1-induced cleavage of mRNA during CrPV and HCV IRES-mediated initiation. (A) Schematic representation of the CrPV IRES with the proximal cognate coding region annotated to show IRES domains 1-3, pseudoknots PKI, PKII and PKIII, stem-loops SL2.1 and SL2.3, inactivating substitutions in domain 2, the P-site codon, the positions of toeprints of IRES-bound 40S subunits and sites of Nsp1-mediated cleavage (red arrows). (B) The influence of *wt* and KH_164-165_AA SARS CoV-2 Nsp1 on association of CrPV IRES mRNA with 40S subunits and 80S ribosomes assayed by toe-printing. (C-E) Cleavage of (C-E) *wt* CrPV IRES mRNA or (E) inactivated SL2.3 and PKIII mutant CrPV IRES mRNA, depending on the presence of *wt* SARS CoV-2 Nsp1 and translational components as indicated, assayed by toe-printing, or where indicated, by primer extension by reverse transcriptase after phenol extraction of mRNA. Positions of the full-length (f.l.) cDNA, 40S/IRES ribosomal complexes, and cleavage sites (red arrows) are shown on the sides of the panels. (F) Cleavage of *wt* EMCV IRES mRNA depending on the presence of *wt* SARS Cov-2 Nsp1 and translational components as indicated, assayed by primer extension by reverse transcriptase after phenol extraction of mRNA. Positions cleavage sites (red arrows) are shown on the right side of the panel. (G, H) The influence of *wt* SARS CoV-2 Nsp1 on the integrity of the HCV IRES depending on the presence of indicated translation initiation components, assayed by toe-printing, or where indicated, by primer extension by reverse transcriptase after phenol extraction of mRNA. Positions corresponding to the full-length (f.l.) cDNA, 40S/IRES and eIF3/IRES contacts, and 48S complexes are shown on the sides of the panels. Lanes C, T, A, and G show the corresponding sequence derived using the same primer as used for primer extension. Separation of lanes in panel F by a white line indicates that these parts were juxtaposed from the same gel.

Although the CrPV IRES was shown to be able to bind to 40S subunits simultaneously with Nsp1 (Yuan et al., 2020), Nsp1 affected ribosomal interaction and positioning of the IRES (Lokugamage et al., 2012; Yuan et al., 2020). Consistent with the previous report (Lokugamage et al., 2012). pre-incubation of 40S subunits with Nsp1 did not induce cleavage of CrPV mRNA, but in contrast to both reports (Lokugamage et al., 2012; Yuan et al., 2020), it also did not affect the 40S/CrPV IRES interaction and accommodation of mRNA in the mRNA-binding channel, yielding the same intensity toe-prints corresponding to 40S/IRES and 80S/IRES complexes (Figure 3B). The reason for the observed discrepancy could result from differences in the coding regions. In our study, the IRES was followed by a long native viral coding region, whereas in the previous studies, the coding region was either native but short (Yuan et al., 2020) or had been replaced by a heterologous reporter (Lokugamage et al., 2012). Although it was possible that binding of the IRES to 40S/Nsp1 complexes displaced Nsp1 from the 40S subunit in our experiments, another study had previously shown that after binding of 40S/Nsp1 complexes to CrPV IRES mRNA containing a 48nt-long coding region, Nsp1 remained long associated with the 40S subunit complex (Lapointe et al., 2021). This prompted us to investigate the ability of different eIFs to support Nsp1-induced cleavage of CrPV IRES mRNA.

Although the CrPV IRES and eIF2/Met-tRNA_iMet_ would clash on the 40S subunit, the simultaneous presence of the IRES and eIF3 is allowed (Pestova et al., 2004). Strikingly, inclusion of eIF3 in reaction mixtures resulted in strong Nsp1-induced cleavage at nt. 6246-6247, 18 nt. downstream from the toe-print corresponding to the 40S/IRES complex, and an additional low-intensity cleavage at nt. 6256 (Figure 3C, lanes 5 and 12). Additional inclusion of eIF4A/eIF4G resulted in 3 prominent discrete downstream cleavages, again ∼6-9 nt. apart (Figure 3C, lanes 4 and 11; summarized in Figure 3A). Further supplementation of reaction mixtures with eIFs 2, 1, 1A and Met-tRNA_iMet_ enhanced accumulation of short mRNA products (Figure 3C, lanes 6 and 13). eIF4A/eIF4G could potentially stimulate cleavage at the first 5’-terminal site and promote downstream cleavage without prior dissociation of truncated mRNA products. Alternatively, downstream cleavage of truncated mRNA products could result from their de novo 5’end-dependent ribosomal attachment to 40S subunits associated with various sets of eIFs like in the case of (CAA)_4_-MF β-globin mRNA. The previously reported lack of cleavage of CrPV IRES mRNA in rabbit reticulocyte lysate upon addition of SARS CoV Nsp1 (Kamitani et al., 2009) could be due to initiation occurring by direct binding of the IRES to 80S ribosomes rather than to 40S subunits because of their engagement into 43S complexes (Pestova et al., 2004).

Next, we determined whether eIF3 can be replaced by any of its subunits, focusing on the eIF3b-g-i module which, like Nsp1, resides at the mRNA entrance (des Georges et al., 2015). We found that the individual recombinant eIF3g subunit was able to substitute for native eIF3 in supporting Nsp1-induced cleavage (Figure 3D, lanes 6 and 7). However, in contrast to native eIF3, inclusion of eIF4A/eIF4G in reaction mixtures containing eIF3g did not stimulate downstream cleavages (Figure 3D, lanes 9 and 10), which is consistent with the lack of interaction between eIF3g and eIF4G and therefore the lack of coupling of the eIF4A/eIF4G helicase activity with ribosomal complexes. Importantly, cleavage was IRES-dependent, and mutations in the 40S-interacting SL2.3 loop and destabilizing mutations in PKIII (Figure 3A), which strongly impair 40S/IRES complex formation (Costantino and Kieft, 2005), also impaired Nsp1-induced cleavage (Figure 3E, lanes 6-15). eIF3g was not able to replace native eIF3 in supporting Nsp1-induced cleavage on EMCV IRES mRNA, highlighting the requirement for native eIF3 for ribosomal attachment to this IRES (Figure 3F).

We also investigated the influence of Nsp1 on the hepatitis C virus (HCV) IRES. As in the case of the CrPV IRES, ribosomal recruitment to the HCV IRES occurs by its direct binding to the 40S subunit, although further initiation events differ drastically, and initiation requires eIF2/Met-tRNA_iMet_ (Pestova et al., 1998b). Association of the HCV IRES with the 40S subunit also displaces eIF3, usurping its ribosomal contacts (Hashem et al., 2013b). In 40S/IRES binary complexes, Nsp1 eliminated toe-prints at nt.355-358 that signal correct accommodation of mRNA in the mRNA-binding channel but did not affect toe-prints at nt.319-312 and nt.342-345 that correspond to upstream 40S/IRES contacts (Figures 3G-H, lanes 1-3). It strongly reduced toe-prints corresponding to 48S complexes (Figure 3G, lanes 4-5; Figure 3H, lanes 6-7). Addition of eIF3 did not stimulate cleavage (Figure 3G, lanes 6-9; Figure 3H, lanes 4 and 10). Although this could be explained by IRES-mediated displacement of eIF3 from the 40S subunit, individual eIF3g also did not support cleavage (Figure 3H, lanes 5 and 11). Thus, in contrast to the CrPV IRES, in Nsp1-containing ribosomal complexes, the coding region of the HCV IRES mRNA is not accommodated in the mRNA-binding channel and therefore does not reach the nuclease active center.

In conclusion, reconstitution of Nsp1-induced cleavage of the CrPV IRES mRNA revealed that it required the minimal set of translational components that consisted of only a 40S subunit and eIF3g.

### Mutational analysis of eIF3g

The 320aa-long eIF3g comprises the N-terminal region, which is involved in formation of the eIF3b-g-i module (Asano et al., 1998; Herrmannová et al., 2012), and the C-terminal RRM domain (aa 232-320) (Cuchalová et al., 2010; Brito Querido et al., 2020). The eIF3b-g-i module interacts with the 40S subunit through the contacts of the ϕ3-propeller domain of eIF3b with the ribosomal protein uS4 (des Georges et al., 2015; Brito Querido et al., 2020), whereas the C-terminal RRM domain of eIF3g was recently assigned to cryo-EM density at the mRNA entry, and accordingly, interacts with helix (h) 16 of 18S rRNA and ribosomal proteins uS3 and eS10 (Brito Querido et al., 2020). To determine the region of eIF3g required for Nsp1-mediated mRNA cleavage, we expressed a series of deletion mutants guided by its secondary and tertiary structure elements (Figure 4A). The eIF3g RRM domain was necessary and sufficient for Nsp1-induced cleavage of the CrPV IRES mRNA (Figure 4B).

**Figure 4.**
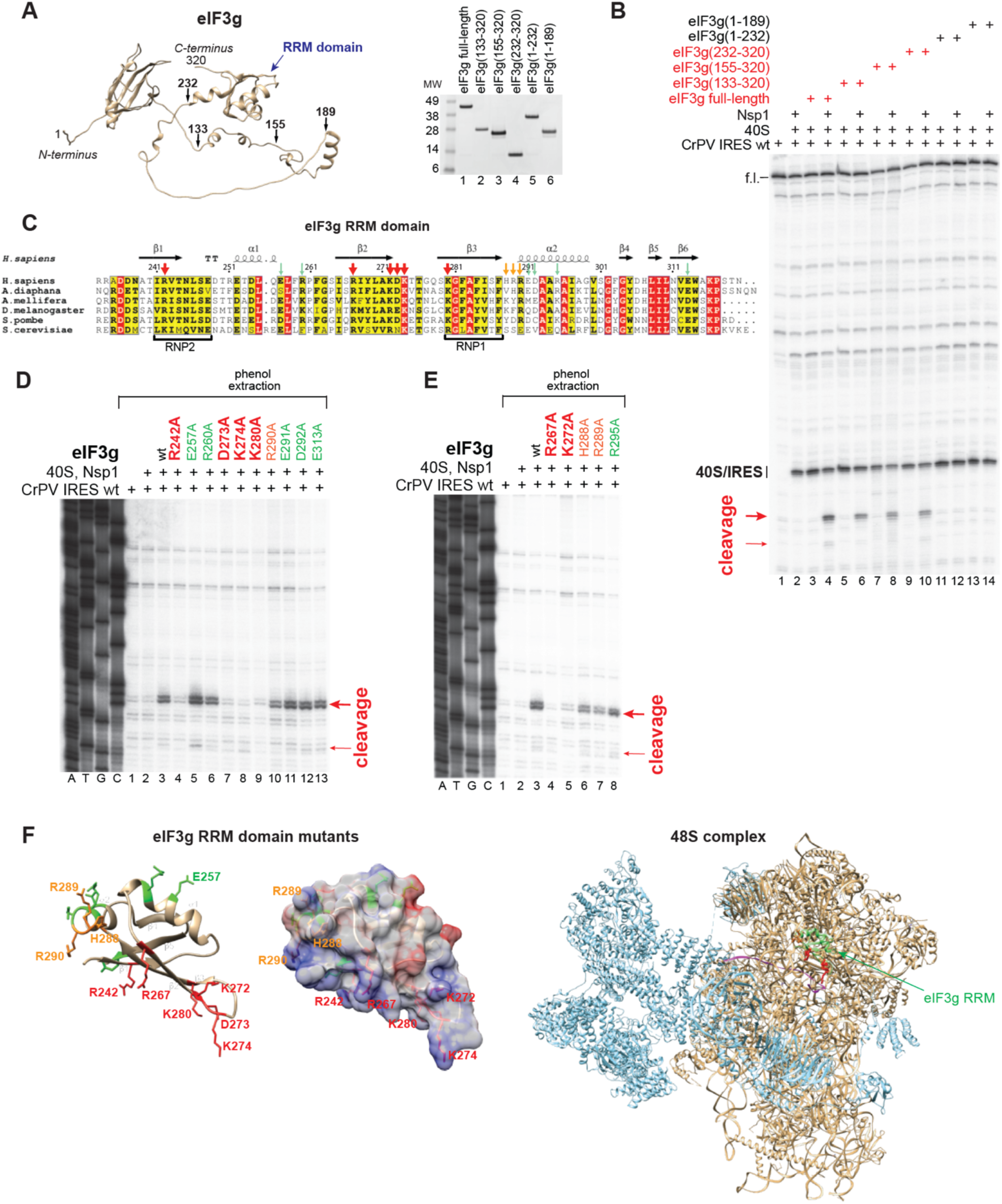
Determination of eIF3g regions required for Nsp1-induced cleavage of CrPV IRES mRNA. (A) Left panel - model of the Alphafold structure of human eIF3g labeled to show the RRM domain and amino acid residues at the borders of eIF3g truncation mutants (black arrows); right panel - recombinant purified eIF3g truncation mutants, assayed by SDS-PAGE followed by SimplyBlue staining. (B) The activities of eIF3g truncation mutants (panel A) in supporting cleavage of *wt* CrPV IRES mRNA induced by *wt* SARS CoV-2 Nsp1, assayed by toe-printing. Positions of the full-length (f.l.) cDNA, 40S/IRES ribosomal complexes, and cleavage sites (red arrows) are shown on the left. (C) Amino acid sequence alignment of the RRM domain of human eIF3g with that of other species, done using Clustal Omega (Sievers et al., 2011). The figure was generated using ESPript 3.0 (Robert and Gouet, 2014). Residues boxed in red are identical. Secondary structure elements from the structure of the human eIF3g RRM domain (PDB: 2CQ0) are represented schematically above and the RNP1 and RNP2 motifs are indicated below the sequence alignment. The amino acid sequence of *H. sapiens* eIF3g (NP_003746.2) between residues 233 and 320 is aligned with its *Exaiptasia diaphana* (XP_020896357), *Drosophila melanogaster* (NP_650887), *Apis mellifera* (XP_391934), *Schizosaccharomyces pombe* (NP_595727) and *Saccharomyces cerevisiae* (NP_010717) homologs. Residues substituted by Ala are indicated by arrows - mutations inactivating Nsp1-induced cleavage are in red, mutations causing a minor reduction in the efficiency of cleavage are in orange, mutations that had no effect on cleavage are in green (summary of results presented in panels D-E). (D, E) The activities of eIF3g RRM domain Ala substitution mutants in supporting cleavage of *wt* CrPV IRES mRNA induced by *wt* SARS CoV-2 Nsp1, assayed by primer extension by reverse transcriptase after phenol extraction of mRNA. Positions of cleavage sites (red arrows) are shown on the right. Mutations inactivating Nsp1-induced cleavage are in red, mutations causing a minor reduction in the efficiency of cleavage are in orange, mutations that do not affect cleavage are in green. Lanes C, T, A, and G show the corresponding sequence derived using the same primer as used for primer extension. (F) Ribbon diagram (left panel) and surface charge distribution (middle panel) of the eIF3g RRM domain (PDB: 6ZMW) annotated to show mutated residues (sticks) colored accordingly to their activity (panels D, E). Right panel - the 48S initiation complex model (PDB: 6ZMW) showing the RRM domain of eIF3g (green ribbon) with amino acids that are essential for Nsp1-induced mRNA cleavage (red and orange sticks), other subunits of eIF3 (light blue ribbon), 40S ribosomal subunit, other initiation factors and Met-tRNA_iMet_ (tan ribbons) and mRNA (magenta ribbon).

The canonical RRM domain adopts a ϕ31α1ϕ32ϕ33α2ϕ34 fold with an antiparallel four-stranded ϕ3-sheet packed against two perpendicular α-helices. The ϕ3-sheet contains two RNP motifs located in ϕ31 and ϕ33 strands. To examine the functional importance of specific regions of eIF3g in supporting Nsp1-induced mRNA cleavage, we generated eIF3g mutants with Ala substitutions of individual charged surface-exposed residues in the RRM domain guided by the model of the 48S complex (Brito Querido et al., 2020), including residues in both RNP motifs and in the conserved positively-charged loop between β2 and β3 strands, which was modeled to interact with h16 (Figure 4C). The activities of all tested mutants are shown in Figures 4D-E: mutants that lost or had severely reduced activity are in red, whereas mutants with moderately reduced activity are in orange. Substitution of key residues in RNP1 position 1 (K280A) and RNP2 position 2 (R242A) abrogated Nsp1-induced cleavage of the CrPV IRES mRNA (Figure 4D, lanes 4 and 9). Notably, there is a basic amino acid at RNP2 position 2 in all eIF3g RRM domains (Figure 4C), whereas the canonical RNP2 motif contains an aromatic residue in this position (Muto and Yokoyama, 2012). Substitution of K274 in the loop between β2 and β3 and of the neighboring K272 and D273 also abrogated mRNA cleavage (Figure 4D, lanes 7-8; Figure 4E, lane 5). An equally strong inactivating effect was observed after R267A substitution in the β2 strand (Figure 4E, lane 4). A basic amino acid at this position is a conserved feature of eIF3g (Figure 4C). Substitution of the neighboring H288, R289 and R290 caused only a minor impairment of the efficiency of cleavage (Figure 4D, lane 10; Figure 4E, lanes 6-7). Interestingly, we consistently observed slightly elevated cleavage in the presence of eIF3g with the E257A substitution in helix α1 (Figure 4D, lane 5).

Mutations in the eIF3g RRM domain that impaired Nsp1-induced mRNA cleavage included critical residues in its RNP motifs and were located on the surface above the mRNA-binding channel (Figure 4F).

To determine whether eIF3g is equally important for Nsp1-induced cleavage of 5’end-dependent and EMCV IRES-containing mRNAs, which require complete eIF3 for ribosomal attachment, we employed native eIF3 containing either a *wt* or a mutated eIF3g subunit. For this, Expi293 cells were transfected with expression vectors for FLAG-tagged *wt*, R242A or R267A eIF3g. eIF3 was first purified on an anti-FLAG M2 affinity matrix and then by FPLC on monoQ. *Wt* and mutated FLAG-tagged eIF3g efficiently incorporated into eIF3, which contained the canonical set of other subunits (Figure 5A). eIF3 containing *wt*, R242A or R267A FLAG-tagged eIF3g were equally active in supporting 48S complex formation on (CAA)_4_-MF β-globin mRNA (Figure 5B, lanes 2, 4 and 6) and on EMCV IRES mRNA (Figure 5C, lanes 2, 4 and 6). In all cases, Nsp1 was able to inhibit 48S complex formation (Figures 5B-C, lanes 3, 5 and 7). However, eIF3 containing R242A mutant eIF3g was substantially less active than eIF3 containing *wt* eIF3g in supporting Nsp1-induced cleavage on all three tested mRNAs: EMCV IRES-containing, CrPV IRES-containing and (CAA)_4_-MF β-globin mRNAs (Figure 5D). The activity of eIF3 containing eIF3g(R267A) was even lower than that of eIF3 containing eIF4g(R242A) in supporting Nsp1-induced cleavage (Figure 5E).

**Figure 5.**
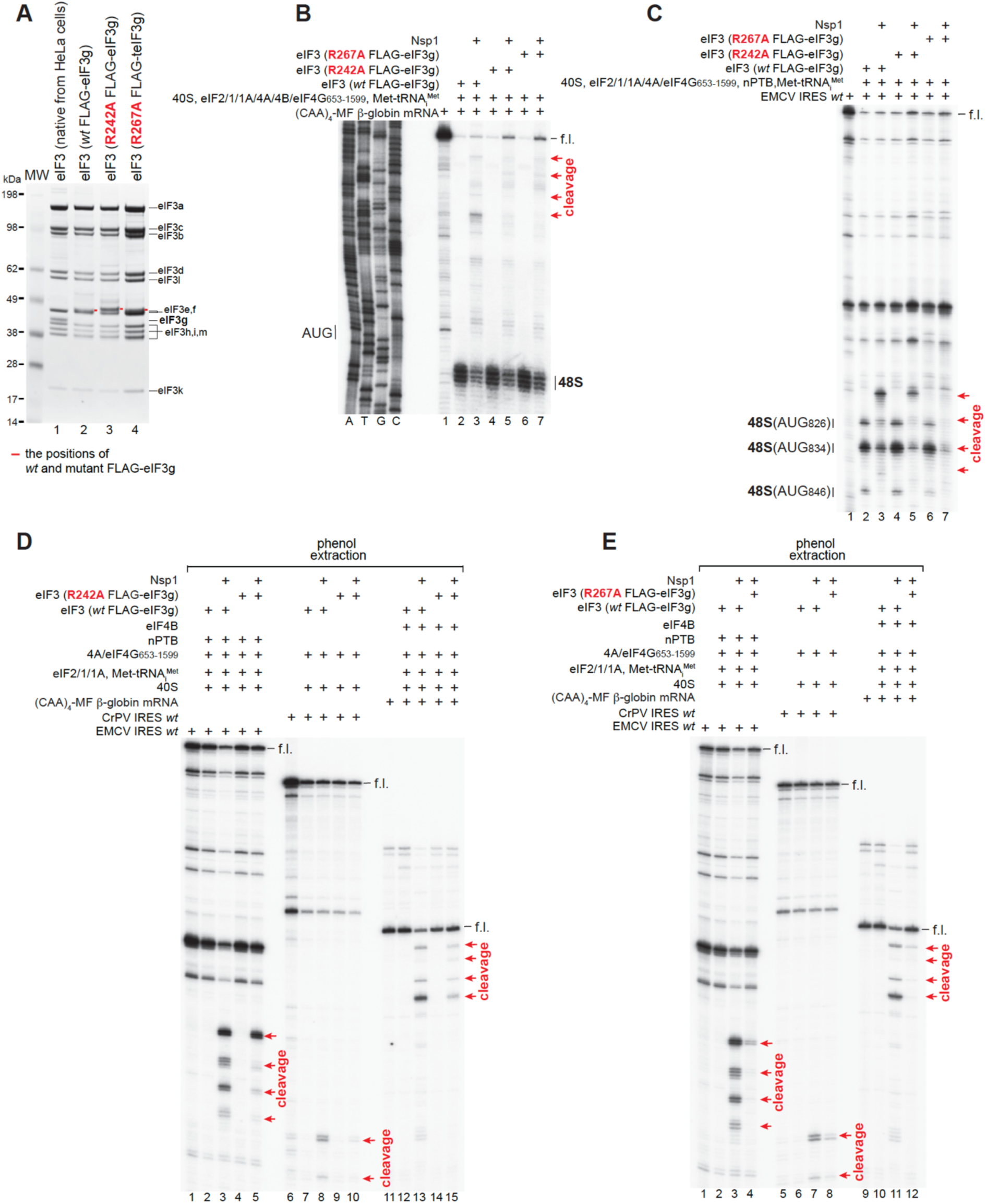
The importance of eIF3g for Nsp1-induced cleavage of β-globin and EMCV IRES-containing mRNAs. (A) Purified native eIF3 from HeLa cells and eIF3 containing FLAG-tagged *wt* or mutant eIF3g expressed in Expi293 cells, analyzed by SDS-PAGE followed by SimplyBlue staining. The positions of individual subunits of eIF3 are indicated. (B-C) The activity of eIF3 containing FLAG-tagged *wt* or mutant eIF3g in 48S complex formation on (B) (CAA)_4_-MF β-globin and (C) EMCV IRES mRNAs in the presence of indicated translational components with/without Nsp1, assayed by toe-printing. (D-E) Nsp1-induced cleavage of (CAA)_4_-MF β-globin, CrPV IRES and EMCV IRES mRNAs depending on eIF3 containing either *wt* or mutant FLAG-tagged eIF3g, assayed by primer extension by reverse transcriptase after phenol extraction of mRNA. (B-E) Positions of the full-length (f.l.) cDNA, 48S complexes and cleavage sites (red arrows) are shown on the sides of the panels.

In conclusion, we determined that during Nsp1-induced cleavage of the CrPV IRES mRNA, the full-length eIF3g could be substituted by its C-terminal RRM domain. Importantly, the RRM domain of eIF3g played an essential role in Nsp1-induced cleavage on all tested mRNAs, irrespective of the modes of their ribosomal attachment.

### Mutational analysis of Nsp1

Next, we performed extensive mutagenesis of the N-terminal domain and the linker region of Nsp1, guided by surface exposure (Clark et al., 2021; Semper et al., 2021), conservation, and charge (Figure S2). As in the case of eIF3g, Nsp1 residues were substituted by Ala. The activities of Nsp1 mutants in inducing mRNA cleavage were essentially the same on different mRNAs irrespective of their mode of initiation. The activities of all tested mutants are shown for the EMCV IRES mRNA in Figures 6A-C (mutants that lost or had severely reduced activity are in red, whereas mutants with moderately reduced activity are in orange). The activities of some selected Nsp1 mutants on CrPV IRES and (CAA)_4_-MF β-globin mRNAs are shown in Figures 6D and 6E, respectively. Mutants with reduced activity preferentially induced cleavage at the 5’-terminal site on the EMCV IRES mRNA indicating the highest affinity of the nuclease to it, which was proportional to the relative loss of the activity (e.g. K11A or R77A versus K129A or H134 Nsp1 mutants) (Figure 6A, lanes 3 and 10; Figure 6B, lanes 10-11). Importantly, in toe-printing experiments done without prior phenol extraction, all inactive/low-activity Nsp1 mutants were able to inhibit 48S complex formation on the EMCV IRES (Figures 6F-H), confirming that they retained the ability to bind to 40S subunits.

**Figure 6.**
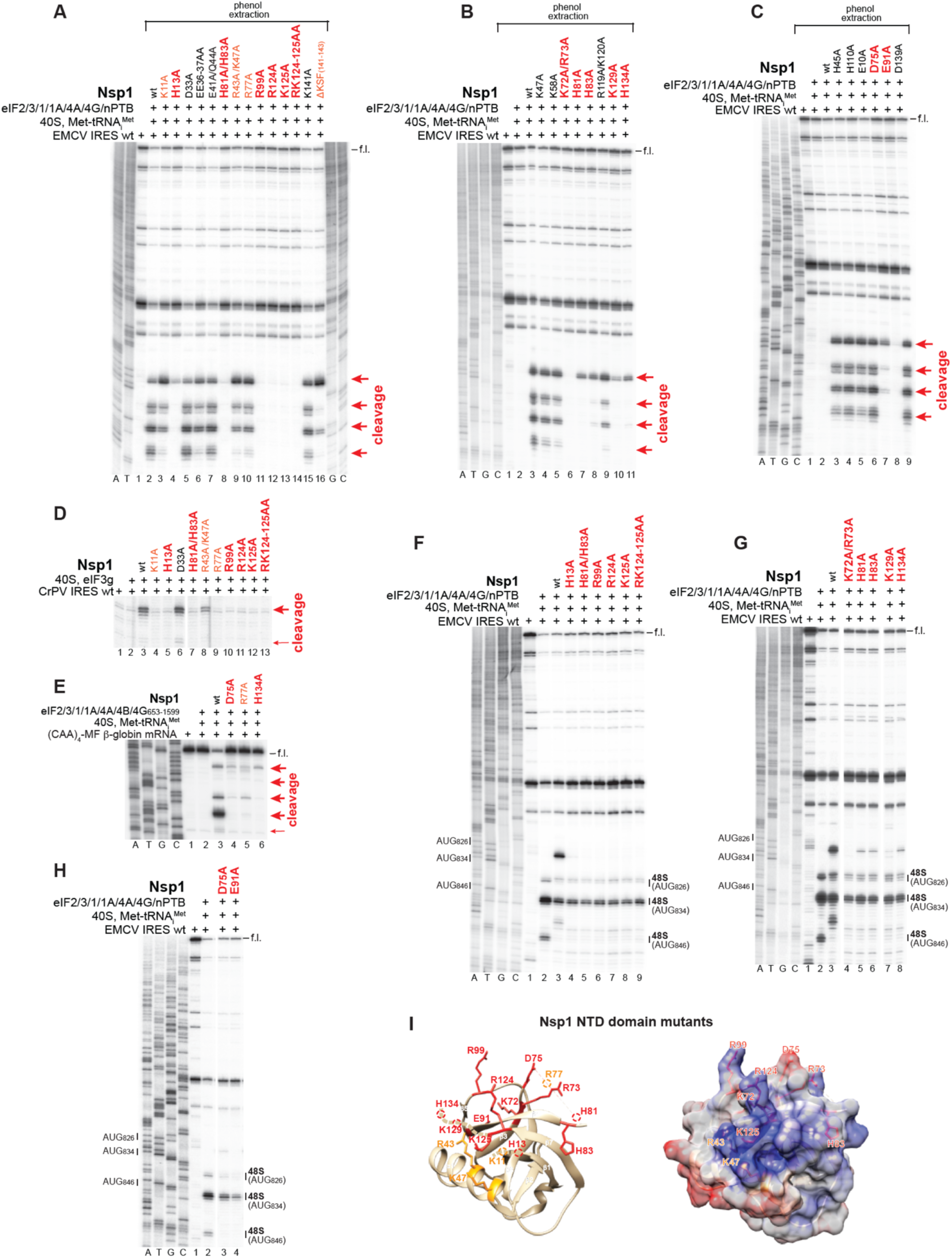
Mutational analysis of SARS CoV-2 Nsp1. (A-E) Cleavage of (A-C) EMCV IRES, (D) CrPV IRES or (E) (CAA)_4_-MF β-globin mRNAs in the presence of *wt* and Ala substitution mutant SARS CoV-2 Nsp1 and indicated translational components, assayed by toe-printing, or where indicated, by primer extension by reverse transcriptase after phenol extraction of mRNA. Mutations having the strongest effect on Nsp1-induced cleavage in panels A-C are in red, mutations causing a moderate effect are in orange, and mutations that don’t affect cleavage are in black. Positions of the full-length (f.l.) cDNA and cleavage sites (red arrows) are shown on the sides of the panels. Lanes C, T, A, and G show the corresponding sequence derived using the same primer as used for primer extension. (F-H) Inhibition of 48S complex formation on the EMCV IRES by Nsp1 Ala substitution mutants with the lowest cleavage-inducing activity, assayed by toe-printing. Positions of the full-length (f.l.) cDNA and 48S complexes are shown on the sides of the panels. Lanes C, T, A, and G show the corresponding sequence derived using the same primer as used for primer extension. Separation of lanes in panels D, G and H by white lines indicates that these parts were juxtaposed from the same gels. (I) Ribbon diagram (left panel) and surface charge distribution (right panel) of the Nsp1 N-terminal domain (PDB: 7K3N) in (left panel) ribbon and (right panel) space-filling representations, annotated to show residues in red or orange sticks that contribute strongly or moderately to Nsp1-induced cleavage, respectively (panels A-C).

Preferential 5’-terminal cleavage of (CAA)_4_-MF β-globin mRNA by impaired Nsp1 mutants (Figure 6E) was consistent with time course data showing that cleavage is sequential and starts from the 5’-terminal site (Figure 1F). Mapping of the residues important for mRNA cleavage onto the crystal structure of the N-terminal domain of Nsp1 showed that they formed an essential surface that was mostly positively charged (Figure 6I). Overall, the essential residues could potentially be involved in the interaction with mRNA, translational components (40S subunits and/or eIF3g) or catalysis of mRNA cleavage *per se*. Taking into account amino acid residues that have established catalytic functions in endonucleases (Yang, 2011), we focused on mutating conserved His and Asp/Glu residues that might contribute to Nsp1-induced catalysis. Abrogation of cleavage was observed in the case of the E91A substitution (Figure 6C, lane 8), whereas H13A (Figure 6A, lane 4), H81A, H83A, H134A (Figure 6B, lanes 7, 8 and 11) and D75A (Figure 6C, lane 7) substitutions very strongly inhibited but did not abolish Nsp1-induced cleavage.

### Inhibition of Nsp1-induced cleavage

Nsp1 is a potential target for inhibition because of its importance for pathogenesis and its sequence and structural conservation. Several chemicals and FDA-approved drugs that could be modified or repurposed to serve as therapeutic SARS CoV-2 inhibitors have been validated by analysis of binding to Nsp1 (Afsar et al., 2022; Borsatto et al., 2022; Kumar et al., 2022; Ma et al., 2022) or by inhibition of its function in cells (Afsar et al., 2022; Kao et al., 2022). Candidate inhibitors include (a) montelukast, a leukotriene receptor antagonist that requires the Nsp1 C-terminal region for binding, modestly impaired SARS CoV-2 replication and inhibited Nsp1-mediated cytopathic effects (CPE) (Afsar et al., 2022; Kumar et al., 2021; Kao et al., 2022), (b) artesunate, a derivative of an antimalarial drug that may disrupt Nsp1 structure (Gurung et al., 2022), (c, d) the tyrosine kinase inhibitor Ponatinib and the 21-aminosteroid lipid peroxidation inhibitor Tirilazad, which synergize with montelukast in inhibiting Nsp1-mediated CPE (Kao et al., 2022), (e) glycyrrhizic acid (Sharma et al., 2020; Vankadri et al., 2020) and (f) mitoxantrone dihydrochloride, an anthracenedione that binds to the Nsp1 CTD (Kumar et al., 2022).

We tested the influence of these compounds using CrPV IRES mRNA, which requires a minimal set of translational components for Nsp1-mediated cleavage. Only mitoxantrone was active. At 10 μM, it abrogated mRNA cleavage while allowing 40S/IRES complex formation, whereas at 500 μM it also inhibited reverse transcription (Figure 7A, lanes 7-8). Mitoxantrone has a planar heterocyclic ring structure with keto groups at the 9^th^ and 10^th^ positions, hydroxy substituents at the 5^th^ and 8^th^ positions and (hydroxyethylamino)-ethylamino side chains at the 1^st^ and 4^th^ positions (Figure 7B). It is FDA-approved for the treatment of multiple sclerosis and several types of cancer. Its best-characterized activity is as a topoisomerase II inhibitor and DNA intercalator (Evsion et al., 2016), but it also binds to several other proteins (e.g. (Wan et al., 2013). Although it completely inhibited cleavage at 10 μM, Nsp1 activity was progressively reduced starting from 5 μM (Figure 7C). Mitoxantrone’s influence on Nsp1-mediated inhibition of 48S complex formation was assayed using HCV IRES mRNA, on which Nsp1 does not induce cleavage. Mitoxantrone did not impair the ability of Nsp1 to inhibit initiation on this IRES (Figure 7D, lanes 3, 5, 7, and 9) but affected 48S complex formation even in the absence of Nsp1 (Figure 7D, compare lanes 2, 4, 6 and 8). The inhibitory effect of mitoxantrone on 48S complex formation was more pronounced on (CAA)_4_-MF β-globin mRNA (Figure 7E, lanes 2 and 4).

**Figure 7.**
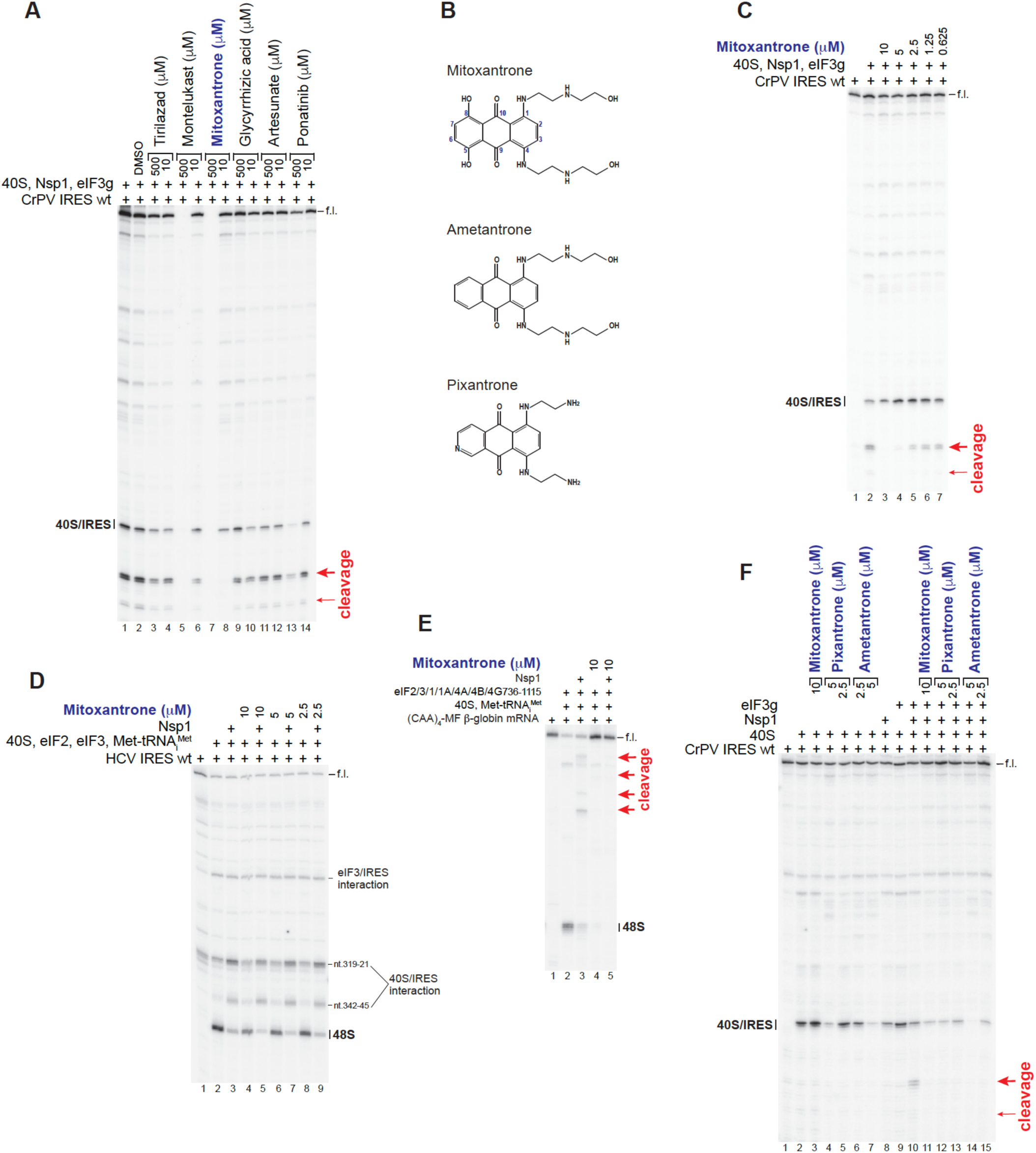
Chemical inhibition of SARS CoV-2 Nsp1-induced cleavage of mRNA. (A, C-F) The influence of FDA-approved drugs on Nsp1-induced mRNA cleavage and inhibition of 48S complex formation on (A, C, F) CrPV IRES, (D) HCV IRES and (E) (CAA)_4_-MF β-globin mRNAs assayed by toe-printing. Positions corresponding to the full-length (f.l.) cDNA, 40S/IRES and eIF3/IRES contacts, 48S complexes and cleavage sites (red arrows) are shown on the sides of the panels. (B) Structures of Mitoxantrone, Ametantrone and Pixantrone.

We also evaluated two related anthracenediones, ametantrone and pixantrone (Figure 7B), to gain insight into the contribution of structural elements to inhibition of Nsp1. Ametantrone lacks hydroxy substituents at the 5^th^ and 8^th^ positions (Zee-Cheng and Cheng, 1978); pixantrone also lacks these groups but has a nitrogen heteroatom inserted into the same ring and ethylamino-diethylamino side chains at the 1^st^ and 4^th^ positions (Menna et al., 2016). Both compounds were even more potent inhibitors than mitoxantrone, abrogating Nsp1-induced cleavage of CrPV mRNA at 2.5 μM (Figure 7F, lanes 13 and 15).

In conclusion, the activity of mitoxantrone showed that Nsp1-induced mRNA cleavage can be inhibited specifically while preserving its ability to inhibit 48S complex formation.

## DISCUSSION

To investigate the mechanism of SARS CoV-2 Nsp1-induced endonucleolytic cleavage of mRNA, we reconstituted this process *in vitro* on three mRNAs (β-globin, EMCV IRES and CrPV IRES mRNAs) that use unrelated mechanisms to initiate translation. Our results argue against the involvement of a putative, as yet-unidentified cellular RNA endonuclease, and establish the essential roles of the C-terminal RRM domain of the eIF3g subunit of eIF3 and Nsp1’s N-terminal domain (NTD) in the cleavage process irrespective of the mode of translation initiation. Requirements for other initiation factors differed for these three mRNAs, reflecting the distinct initiation mechanisms that they use, and specifically the different factor requirements of each for ribosomal attachment.

The Nsp1-induced cleavage of CrPV IRES mRNA required a minimal set of translational components consisting of only a 40S subunit and eIF3g’s RRM domain. The cleavage site was located 18 nts downstream from the toe-prints corresponding to 40S/IRES complexes. The position of the cleavage site and the ribosomal location of eIF3g’s RRM domain at the mRNA entrance (Brito Querido et al., 2020) indicate that Nsp1-induced cleavage takes place on the solvent side of the 40S subunit downstream from the mRNA entrance. Mutational analysis identified a positively charged surface on the Nsp1 NTD and a surface above the mRNA-binding channel on the RRM domain of eIF3g that contain residues essential for cleavage. The key questions concerning the mechanism of cleavage include the exact identity of the nuclease and the individual roles of all components (i.e. the Nsp1 NTD, eIF3g’s RRM domain and the 40S subunit) in this process. A triple-Ala substitution of the critical residues in the RNP motifs of the RRM domain of *S. cerevisiae* Tif35/eIF3g reduces the processivity of scanning and resumption of scanning by post-termination ribosomes (Cuchalová et al., 2010), suggesting that the RRM domain might interact with the backbone of mRNA outside the entry channel. The key residues in the RNP motifs of eIF3g (i.e. R242 and K280) were also important for Nsp1-mediated cleavage, as were other residues in the same cavity that faces the mRNA-binding channel. Moreover, crosslinking-MS analysis of DSSO cross-linked HEK293T cells overexpressing FLAG-tagged Nsp1 revealed extensive cross-links between the Nsp1 NTD and the residues of the eIF3g RRM domain that are surface-exposed in the 43S complex, indicating that the interaction between Nsp1 and eIF3g, either direct or mRNA-dependent, can occur in the context of 43S ribosomal complexes (Graziadei et al., 2022). The cross-linked Lys residues in the Nsp1 NTD (Graziadei et al., 2022) were located on the same surface or in close proximity to residues that were essential for Nsp1-induced cleavage. Thus, eIF3g’s RRM domain could potentially function to correctly orient the Nsp1 NTD, to act as a conduit for mRNA, or even to contribute to catalysis if the nuclease catalytic site is formed not by Nsp1 alone but is a composite of Nsp1 and the RRM domain of eIF3g. Consistently, there are also several not mutually exclusive possibilities for the role of the essential Nsp1 NTD surface: (i) to interact with eIF3g, (ii) to interact with the 40S subunit, and (iii) to interact with mRNA and to form the nuclease active center. The role of the 40S subunit is likely limited to being a scaffold for the spatial arrangement of the other components.

The Nsp1 NTD and the RRM domain of eIF3g also had essential roles in Nsp1-induced cleavage of the 5’end-dependent β-globin and EMCV IRES-containing mRNAs, but in contrast to the CrPV IRES mRNA, cleavage on these mRNAs required additional initiation factors. Thus, cleavage of β-globin mRNA needed group 4 eIFs and intact native eIF3 that are responsible for ribosomal attachment during 5’end-dependent initiation, whereas cleavage of the EMCV IRES mRNA additionally required the eIF2•GTP/Met-tRNA_iMet_ ternary complex. Ribosomal attachment to the EMCV IRES depends on the specific interaction of its JK domain with the central eIF4A-binding domain of eIF4G (Pestova et al., 1996). However, the JK domain is not sufficient for the ribosomal attachment to the IRES, and the requirement of the eIF2 ternary complex for Nsp1-mediated cleavage of the EMCV IRES mRNA suggests the essential, as yet unidentified role of eIF2•GTP/Met-tRNA_iMet_ in ribosomal attachment to the IRES. In all cases, several cleavage sites separated by 6-9 nts were observed.

The time course of Nsp1-induced cleavage of β-globin mRNA indicates that cleavage is sequential rather than random and starts from the 5’end. Consistently, if cleavage was inefficient e.g. as in the presence of some Nsp1 mutants, it did not have time/opportunity to progress further than the 5’-terminal sites. The question remains whether mRNA dissociates from the 40S subunit after the first cleavage and then rebinds, leading to cleavage at the next downstream site as a result of a separate attachment event, or if mRNA remains associated with the 40S ribosomal complexes and group 4 eIFs continuously feed it into the nuclease active center. The 6-9 nt spacing between cleavage sites suggests that cleavage requires the accommodation of the upstream 5’-terminal 6-9 mRNA nucleotides in the nuclease active center. The minor variation in the distance between cleavage sites may be indicative of a small degree of nucleotide specificity of cleavage. Notably, the 5’-terminal cleavage site was located close to the 5’-end of mRNA, and even in conditions of eIF4E-cap interaction during cleavage of native capped β-globin mRNA in the presence of native eIF4F, the first cleavage site was only 7 nt from the cap. Regarding the ribosomal position of the eIF3g RRM domain and its cross-linking to the Nsp1 NTD, it is difficult to reconcile the position of the 5’-terminal cleavage site with the suggested ribosomal position of eIF4E and the proposed slotting mechanism of ribosomal attachment during 5’end-dependent initiation (Brito Querido et al., 2020), because in this case the 5’-terminal mRNA region would be too far from the hypothetical nuclease active center. In contrast to CrPV mRNA, no cleavage in the coding region was observed on β-globin mRNA. Strong inhibition of 48S complex formation on β-globin mRNA by *wt* and mutant Nsp1 indicates that the co-existence of Nsp1 with accommodated mRNA is unique to the CrPV IRES, which does not require the eIF2•GTP/Met-tRNA_iMet_ ternary complex and binds to the A site of the 40S subunit.

In the case of the EMCV IRES mRNA, cleavage occurs in regions that are very close to domain L, and during initiation would most likely be slotted into the mRNA-binding channel. In the presence of Nsp1, this region likely remains on the solvent side of the 40S subunit. Sequence and/or structural features of the region enable two particular areas in it to be independently accommodated in the nuclease active center yielding cleavages at nt. 830-831 and 849-850, respectively. Subsequent downstream cleavage could potentially result from the 5’end-dependent attachment of truncated mRNAs to mRNA-free 43S complexes. The mechanistic basis for downstream cleavages on the CrPV IRES mRNA is also not clear and they could also result from 5’end-dependent attachment of truncated mRNAs to mRNA-free 40S ribosomal complexes.

In conclusion, we identified the essential role of the Nsp1 NTD and the RRM domain of eIF3g in Nsp1-medited cleavage of mRNAs and determined the initiation factor requirements for this process on mRNAs translated by three different initiation mechanisms, thus providing the foundation and the framework for the future structural studies of the molecular mechanism of cleavage.

## MATERIALS AND METHODS

### Construction of plasmids

Vectors for bacterial expression of eIF1, eIF1A, eIF4A, eIF4B, eIF5, eIF4G_736-1115,_ eIF4G_653-1599,_ *E. coli* methionyl tRNA synthetase, nPTB and DHX29 have been described (Pestova et al., 1996; Pestova et al., 1998a; Pestova et al., 2000; Lomakin et al., 2000, 2006; Pilipenko et al., 2001; Sweeney et al., 2021).

The vectors pET15b-His_6_-Nsp1(SARS CoV2) and pET15b-His_6_-Nsp1(SARS CoV) for expression in *E. coli* of Nsp1 from SARS CoV2 (NCBI Ref. seq. YP_009725297.1) and SARS (Urbani strain) (Genbank Acc. No. AAP13442.1) with N-terminal His_6_ tags were made by inserting DNA between NcoI and BamHI sites of pET15b (GenScript, Piscataway, NJ). pET15b-His_6_-Nsp1(SARS CoV2) was then used by GenScript to generate the substitution mutants described in the text.

The vector pET15b-His_6_-eIF3g for expression in *E. coli* of human eIF3g (Genbank Acc. No. NM_003755) with an N-terminal His_6_ tag was made by inserting DNA between NdeI and XhoI sites of pET15b (GenScript). pET15b-His_6_-eIF3g was used by GenScript to generate the deletion and substitution mutants.

Vectors for expression in mammalian cells of N-terminally 3XFLAG-tagged wild type and mutant forms of human eIF3G were prepared by inserting the appropriate wild type ORF into pcDNA3.1(+)-N-DYK and using the resulting vector for mutagenesis (GenScript).

Vectors for transcription of *wt* CrPV IRES, *wt* HCV IRES and (CAA)_4_-MF β-globin mRNAs, tRNA_iMet_ and tRNA^Phe^ have been described (Pisarev et al., 2007; Hashem et al., 2013b; Zinoviev et al., 2018; Abaeva et al., 2020).

pUC57-based vectors for transcription of CrPV IRES nt. 5997-6320 variants with CAGC_6142-45_AAAG substitutions in SL2.3 and CCUA_6134-37_GGAU substitutions in PKIII were made by Synbio Technologies (Monmouth Junction, NJ).

The vectors pUC57-[EMCV nt.373-1656] for transcription of *wt* and mutant EMCV IRES-containing mRNA were made by GenScript and contained a T7 promoter followed by EMCV nt.373-1656, with or without mutations as indicated in the text.

Destabilization (destab. L domain) and stabilization (stab. L domain) of L domain were achieved by CGUGGUUU _802-809_GCACCAAG and AA_791-792_CC/UUU_807-809_GGC substitutions, respectively. L domain deletion mutants were obtained by deletion of nucleotides 790-809 (ΔL domain) or by their substitution by GU_791-792_ (ΔL domain + 2nt), GUUU_791-794_ (ΔL domain + 4nt), and GUUCCUUU_791-798_ (ΔL domain + 8nt). Mutations in the I domain comprised CAG_529-531_GUC substitutions. Mutations in the region surrounding the two 5’-terminal Nsp1 cleavage sites involved G_816_C, G_828_C and G_836_C substitutions. Mutations surrounding 3’-terminal cleavage sites involved deletion of AUG_846_ and introduction of AUG_853_ by G_846_C and A_854_U substitutions.

### Purification of ribosomal subunits, initiation and elongation factors, aminoacyl-tRNA synthetases and recombinant Nsp1

Native mammalian 40S and 60S ribosomal subunits, eIF2, eIF3, eIF4F, eIF5B, eEF1H, eEF2 and aminoacyl-tRNA synthetases were purified from rabbit reticulocyte lysate (RRL) (Green Hectares, Oregon, WI) as described (Pisarev et al., 2007; Zinoviev et al., 2019, 2020). Human recombinant eIF1, eIF1A, eIF4A, eIF4B, eIF5, eIF4G_736-1115_, eIF4G_736-1599_, nPTB and DHX29, and *E. coli* methionyl-tRNA synthetase were expressed in *E. coli* BL21(DE3) (Invitrogen) and purified as described (Pestova et a., 1996; Pestova et al., 1998; Pestova et al., 2000; Lomakin et al., 2000, 2006; Pilipenko et al., 2001; Sweeney et al., 2021).

Recombinant *wt* and mutant forms of Nsp1 and eIF3g were expressed in 2L of *E. coli* BL21 (DE3) (Invitrogen) for 18-20 h at 16°C after induction with 1 mM IPTG, and then purified by affinity chromatography on Ni-NTA-agarose (QIAGEN). *Wt* and mutant Nsp1 were further purified on a FPLC MonoQ HR 5/5 column with fractions being collected across a linear 100-500 mM KCl gradient and eluted at ∼260 mM KCl. *Wt* eIF3g, eIF3g(1-189) and eIF3g(1-232) were also further purified on a FPLC MonoQ HR 5/5 column with fractions being collected across a linear 100-500 mM KCl gradient. Wt eIF3G and eIF3g(1-232) eluted at ∼175 mM KCl, whereas eIF3g(1-189) eluted at ∼320 mM KCl. N-terminally truncated eIF3g(133-320), eIF3g(155-320) and eIF3g(232-320) were purified further on a FPLC MonoS HR 5/5 column with fractions being collected across a linear 100-500 mM KCl gradient and eluted at ∼240, ∼270 and ∼340 mM KCl. The identity of all truncated forms of eIF3g was confirmed by western blotting using antibodies against His_6_-tag (Abcam).

To obtain native eIF3 containing mutant eIF3g subunits, Expi293F cells (Thermo Scientific) were transfected with expression vectors for 3xFLAG-tagged *wt* eIF3g or its mutated forms using Expifectamine 293 according to the manufacturer’s protocol (Thermo Scientific). Cells expressing 3xFLAG-tagged eIF3g (200 ml suspension) were grown for 72 hours, harvested, washed with PBS buffer, and lysed in buffer A (20 mM Tris pH 7.5, 2 mM DTT, 100 mM KCl) using a Dounce homogenizer. Cell debris was removed by centrifugation for 30 min at 45,000 rpm at 4°C using a Beckmann 50.2 Ti rotor, and the supernatant was loaded onto anti-DYKDDDDK G1 affinity resin (GenScript) equilibrated in buffer A, which was subsequently washed with the same buffer. Proteins were eluted by incubation with buffer A containing 0.5 mg/ml 3xFLAG peptide (ApexBio, Houston TX) at room temperature for 30 minutes. The eluted eIF3 was purified further by FPLC on a MonoQ 5/50 GL column: fractions were collected across a linear 100-500 mM KCl gradient. eIF3 containing FLAG-tagged eIF3g eluted at around 300 mM KCl.

Native total calf liver tRNA (Promega) and *in vitro* transcribed tRNA_iMet_ were aminoacylated using recombinant *E. coli* methionyl-tRNA synthetase, and *in vitro* transcribed tRNA^Phe^(GAA) was aminoacylated using native aminoacyl-tRNA synthetases (Lomakin et al., 2006; Pisarev et al., 2007).

### *In vitro* reconstitution and analysis of Nsp1-induced cleavages

To reconstitute Nsp1-induced cleavage of mRNA, 100 nM 40S subunits were first preincubated with 500 nM *wt* or mutant Nsp1 for 10 minutes at 37°C in buffer B (20 mM Tris, pH 7.5, 2 mM dithiothreitol, 0.25 mM spermidine, 2.5 mM MgCl_2_) supplemented with 1 mM ATP, 0.5 mM GTP and 1.5 U/μl RiboLock RNAse inhibitor, after which reaction mixtures were supplemented with 40 nM (CAA)_4_-MF β-globin, native β-globin, *wt* or mutants EMCV IRES, *wt* or mutant CrPV IRES, or *wt* HCV IRES mRNAs and indicated combinations of 250 nM eIF2, 150 nM native or mutant eIF3, 500 nM eIF1, 500 nM eIF1A, 500 nM eIF4A, 400-500 nM eIF4G_653-1599_ or eIF4G_736-1115_ (except for the experiment shown in Figure 1D, in which the concentration of eIF4G_736-1115_ was reduced 3-fold), 100 nM eIF4F, 250 nM DHX29, 1 μM *wt* or mutant eIF3g, 250 nM nPTB and 150 nM *in vitro* transcribed Met-tRNA_iMet_ (for (CAA)_4_-MF β-globin, native β-globin and HCV IRES mRNAs) or total native mRNA aminoacylated using recombinant *E. coli* methionyl-tRNA synthetase (for *wt* and mutant EMCV IRES mRNAs). After that the incubation continued for 15 more minutes except for the time course experiments (Figures 1F and S1A) when aliquots were removed from the reaction mixtures at indicated time points. In some instances (where indicated in Figures 1B and 2B), 40S subunits were not preincubated with Nsp1, which was added for an additional 15 minutes after incubation of all indicated translational components. To investigate the influence of U-74389G, Montelukast, Artesunate, Ponatinib, Mitoxantrone, Glycyrrhizic acid, Ametantrone (Cayman Chemical) and Pixantrone maleate (Selleck Chemicals) on Nsp1 activity, each drug was dissolved in DMSO and added to buffer B to reach the desired concentration.

To investigate the influence of Nsp1 on mRNA in assembled 80S initiation complexes and on the ability of 80S complexes to undergo elongation (Figures 1B and 1E), 80S initiation complexes were assembled on (CAA)_4_-MF β-globin by incubating 40 nM mRNA with 100 nM 40S subunits, 250 nM eIF2, 150 nM eIF3, 500 nM eIF1, 500 nM eIF1A, 500 nM eIF4A, 500 nM eIF4G_736-1115_ and 150 nM *in vitro* transcribed Met-tRNA_iMet_ for 15 minutes at 37°C in buffer B supplemented with 1 mM ATP, 0.5 mM GTP and 1.5 U/μl RiboLock RNAse inhibitor, after which reaction mixtures were supplemented with 130 nM 60S subunits, 400 nM eIF5 and 150 nM eIF5B, and incubation continued for another 15 minutes. Assembled 80S initiation complexes were then incubated for 15 minutes with indicated combinations of 500 nM Nsp1, 150 nM eEF1H, 400 nM eEF2 and 150 nM Phe-tRNA^Phe^.

To detect cleavage, mRNAs were analyzed either directly by toe-printing in the presence of translational components and drugs, or by primer extension after phenol-extraction and ethanol precipitation. In all cases, primer extension reactions were done using AMV reverse transcriptase (Promega) and ^32^P-labeled primer (Pisarev et al., 2007). Radiolabeled cDNAs were phenol-extracted, ethanol-precipitated, resolved on 6% polyacrylamide gel and analyzed by phosphoimager.

## COMPETING INTEREST STATEMENT

The authors declare no competing interests.

## ACKNOWLEDGMENTS

We thank Andrew Tcherepanov for expert technical assistance. This work was supported by NIH grants GM122602 to T.V.P, and NIH grant AI166944 to T.V.P. and C.U.T.H.

## AUTHOR CONTRIBUTIONS

I.S.A., Y.A. and A.M. performed all experiments. T.V.P., C.U.T.H., I.S.A., Y.A. and A.M. designed experiments and interpreted data. T.V.P. and C.U.T.H. wrote the paper with input from all authors.

